# Combined topological data analysis and geometric deep learning reveal niches by the quantification of protein binding pockets

**DOI:** 10.1101/2023.08.25.554762

**Authors:** Peiran Jiang, Jose Lugo-Martinez

## Abstract

Protein pockets are essential for many proteins to carry out their functions. Locating and measuring protein pockets as well as studying the anatomy of pockets helps us further understand protein function. Most research studies focus on learning either local or global information from protein structures. However, there is a lack of studies that leverage the power of integrating both local and global representations of these structures. In this work, we combine topological data analysis (TDA) and geometric deep learning (GDL) to analyze the putative protein pockets of enzymes. TDA captures blueprints of the global topological invariant of protein pockets, whereas GDL decomposes the fingerprints to building blocks of these pockets. This integration of local and global views provides a comprehensive and complementary understanding of the protein structural motifs (*niches* for short) within protein pockets. We also analyze the distribution of the building blocks making up the pocket and profile the predictive power of coupling local and global representations for the task of discriminating between enzymes and non-enzymes. We demonstrate that our representation learning framework for macromolecules is particularly useful when the structure is known, and the scenarios heavily rely on local and global information.

## 1 INTRODUCTION

Proteins are biological macromolecules responsible for carrying out many of the essential functions of cells. Understanding protein function remains a fundamental aim to understand life at the molecular level. While the availability of protein sequence and structure information has grown exponentially, the experimental determination of the function of a protein is still limited by time and cost. To address this limitation, a plethora of computational methods that predict protein function have been developed over the years [37]. Key to these computational approaches is to infer protein function by finding proteins with similar sequence, structure, or other characteristics. For instance, the shape and properties of the protein surface determine what interactions are possible with ligands and other macromolecules [13].

Among the multiple elements of protein structures, voids, pockets, and channels are important features of protein surface, thus, play a crucial role for many protein functions [24]. For instance, locating and measuring protein pockets and cavities has been shown to be useful for computer aided drug design [30]. Furthermore, studying the anatomy of protein pockets and cavities with geometry and topology helps us understand the shape and topological information niches, defined as protein structural motifs [32].

Toward this goal, a natural approach is to use topological data analysis (TDA) for capturing the global information within protein structure data [10]. For instance, persistent homology (PH), a main workhorse of TDA, could represent macromolecules into persistent bar codes, diagrams, or landscapes as input to machine learning models [6]. PH can help reduce the structural complexity and preserve the topological invariant properties. The encoded features, including connected components, loops, voids, and other higher-order properties along with their persistence are descriptors of global information in the Euclidean space ℝ ^*n*^. Traditional TDA-based methods are not able to capture the local structural information since topology studies properties of spaces that are invariant under any continuous deformation [15]. For example, PH only captures changes of topological invariants and provides some persistence, which is not sensitive to homotopic shape evolution. Fortunately, past and ongoing research precisely address the shortcoming of TDA when it comes to protein morphology study. Zixuan *et al*. proposed a topological approach for protein classification [7]. Kovacev-Nikolic *et al*. profiled the persistence landscapes of to protein structures.Both work demonstrated that TDA and PH based methods are able to analyze the protein structures. Cases have shown TDA are strong enough to even identify protein superdomains [7] or the patterns of maltose-binding protein [22]. However, according to the continuous effort of enzyme structure-function relationships, we still need more biochemical and biophysical information as analytical tools to better understand enzymes universally[5].

Therefore, we need to consider more detailed features in biochemistry and biophysics towards shapes and heterogeneous properties of structural data. An alternative approach involves the use of computational geometry such as geometric deep learning (GDL). GDL is a locally-aware tool that gives us zoomed-in view of the data and maps it into the representation space with domain knowledge [2]. Additionally, GDL is a technique that generalizes neural networks to non-Euclidean domains such as graphs, manifolds, meshes, among other representations. Unlike traditional computational geometry, GDL can handle information beyond distance, mesh, shape descriptors, or curvature descriptors, including extended geometry-associated features such as node or edge labels as well as different types of interactions to name a few. In the context of proteins, amino acid residues are the building blocks of proteins. These residues can be further categorized by shared biochemical and biophysical properties. Moreover, pockets with similar topology but different microenvironments caused by residue composition (e.g., charged, pH, hydrophilic, hydrophobic) are likely to exhibit distinct binding affinity profiles. Therefore, GDL is a suitable approach to explore these local biochemical features of protein structures and make complementary contributions to those structure profiles obtained via TDA.

Inspire by recent efforts that have shown that global structural topology and local geometry refinement can have mutual benefits towards protein pocket mining [29, 31], we combine TDA and GDL to analyze the putative protein pockets of enzymes. We study the building information of protein pockets via the use of fully labeled hypergraphs [26] and feed them as input to hypergraph neural networks [21]. Given the success of TDA and GDL within the protein space, we hypothesize that the combination of TDA and GDL will reveal niches of protein binding pockets by leveraging the quantitative power of top-down and bottom-up representations. In particular, the contributions of this work are (1) We show that GDL successfully decomposes the blueprints to building blocks with nuanced biochemical features, whereas TDA captures the global topological invariant of pockets in terms of structure. The combination of both views gives us a comprehensive and complementary understanding of the niches within the space of protein pockets. (2) We analyze and predict the propensity that proteins become an enzyme with such pockets. Furthermore, we show that these predictions are supported by enzymology. (3) We construct a novel representation learning framework for proteins. This framework is particularly useful when the structure is known, and the down-stream task heavily relies on local and global information.

## 2 RELATED

### WORK Multiparameter persistent homology

Multiparameter persistent homology (MPH) is an extension to persistent homology of single filtered space. As an active area of TDA, MPH can capture the topological invariants of interest by considering the multi filtered space. MPH provides and calculates (*n*-parameter) persistence modules (algebraic invariants of data), simply by applying homology field coefficients to a multifiltration [4]. MPH gives us insights to interpret and compare data at different types and scales simultaneously. For example, MPH has been used to study immune cell distributions with differing oxygenation levels [34]. In this work, we aim to apply MPH on protein pockets with different pocket confidence levels.

### Hypergraph kernels and hypergraph neural networks

Hypergraphs, a generalization of graphs, provide a flexible and accurate model to encode higher-order relationships inherently found in many disciplines. In particular, *hypergraphlets*, small hypergraphs rooted at a vertex of interest, have been successfully used to probe large hypergraphs as hypergraphs can be thought of as being composed of a collection of independent hypergraphlets [16, 26]. Furthermore, Lugo-Martinez *et al*. [2020] present a generalized algorithm for counting hypergraphlets as means to define a kernel method on vertex and edge-labeled hypergraphs for analysis and learning.

To leverage the expressiveness of hypergraphs, researchers have tried to adapt graph neural network (GNN) into hypergraphs graph neural networks (HGNNs) for graph representation learning. The challenge is how to learn powerful representative embeddings without losing such higher order information. Huang and Yang [2021] proposed a unified framework for graph and hypergraph neural network trying to unify the message passing process with minimal effort. The message passing in hypergraphs is shown to be as powerful as the 1-dimensional Generalized Weisfeiler-Lehman (1-GWL) algorithm in terms of distinguishing non-isomorphic hypergraphs [3].

### Topological layers

Many studies have incorporated topological invariants for end-to-end learning with neural networks. Hofer *et al*. [2019] proposed the first topological layer by using the idea of Gaussian transformation on persistence diagrams. They also proposed a novel type of readout operation to leverage persistent homology computed via a real-valued, learnable, filter function layer [17]. Another more comprehensive layer called *PersLay* for persistence and topological signatures was described by Carriere *et al*. [2020]. PersLay is an end-to-end, differentiable framework for learning versatile PH descriptors in a neural network, which allows us to better understand the topological features of data in an automatic way. Various vectorization methods were used in PersLay for better learnable representations of persistent diagrams. Finally, Horn *et al*. [2021] proposed a topological neural network which is strictly more expressive than message passing GNNs.

## 3 BACKGROUND AND NOTATION

Here we review the background on protein pockets as well as some basic concepts and notations of TDA and GDL.

### Protein binding pocket

As mentioned earlier, protein binding pockets (ligand binding sites or simply pockets) play an important in protein function and computer aided drug design. Pockets are regions with specific sizes, shapes, and physicochemical properties. On the other hand, ligands are specific small molecules that could fit into pockets and bind with host proteins. Different approaches have been utilized to predict ligand binding sites including geometric-, energetic-, consensus-, template-, conservation-, and knowledge-based methods. A comprehensive review of these approaches is provided by Krivák & Hoksza [2018].

For simplicity, in this work, we treat a protein *P* of length *n* as a sequence of amino acid residues denoted as 𝒮= *s*_1_*s*_2_*s*_3_ *s*_*n*_, where each *s*_*i*_ ∈ 𝒮 represent one of the 20 common amino acids (Appendix B). These residues are quite diverse in terms of geometry, charge, hydrophobicity, polarity, and other properties. A protein pocket with *n*_*C*_ amino acids 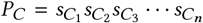 is a special subset of amino acids that are spatially closed within a 3-D structure.

### Vietoris–Rips complex

A simplex is a generalization of the notion of a triangle or tetrahedron to arbitrary dimensions. A *k*-simplex is a *k*-dimensional polytope that is the convex hull of its *k* + 1 vertices. A simplicial complex 𝒦 is a set composed of simplices and satisfies the following conditions: (1) every face of a simplex from 𝒦is also in 𝒦, and (2) the non-empty intersection of any two simplices σ_1_, σ_2_ ∈ 𝒦is a face of both σ_1_ and σ_2_.

A Vietoris–Rips complex consists of all those simplices whose vertices are at pairwise distance no more than *r* as

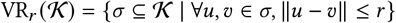

In this work, we only consider the metric space, (*χ, d*_*χ*_), where *χ* is the set of alpha-carbon coordinates of pockets and *d*_*χ*_ is the Euclidean distance between them.

### Multiparameter persistence homology and landscape

Multiparameter persistence homology is an extension of (single-parameter) persistence homology [8, 33]. More formally, MPH is induced by a *multifiltration function f* : *X* → ℝ^*d*^. For any *a, b* ∈ ℝ^*d*^, we denote *a* ≺*b* when ∀*i, a*_*i*_ ≤ *b*_*i*_. Then the sub-level sets *F*_*r*_ = {*x*∈*X* | *f* (*x*) ≤ *r* } satisfy *F*_*a*_ ⊆*F*_*b*_ as long as *a* ≺*b*. For a family of multiparameters *r*_1_, *r*_2_,, *r*_*n* ∈_ ℝ^*d*^, when *r*_*i*_ ≤*r* _*j*_, the sets 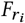 and the inclusion relationships 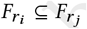 is called a *multifiltration* of *f*. To get the homology, we apply the homology functor *H*_*k*_, which maps topological spaces to vector spaces. *H*_*k*_ (*F*_*r*_) is under-stood as the *k*th topological features of *F*_*r*_. The sequence of vector spaces connected with linear maps (*H*_*k*_ (*F*_*a*_) →*H*_*k*_ (*F*_*b*_)) is called as *persistence module* of *f, M* (*f*). The canonical decomposition of a persistence module is the sum of simple modules.

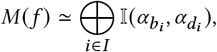

where 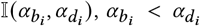 is the *interval module*. An interval module intuitively represents a topological feature that appeared at parameter *α*_*b*_ and disappeared at parameter *α*_*d*_ in the filtration. A representation of decomposition *M*(*f*) in a plane is called the *persistence diagram*. Then the single parameter persistence landscape of *M*(*f*) could be defined as

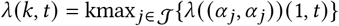

where 𝒥 is the set of associated persistence diagram given by the indexed pair set 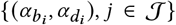. kmax is the *k*th largest value operator of the indexed set. The multiparameter persistence landscape is similarly defined as

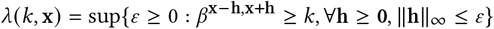

The multiparameter persistence landscape considers the maximal radius over which *k* features persist in every (positive) direction of x [33].

### Fully labeled hypergraphs

A hypergraph *G* has both a vertex set *V* and an edge set *E*, where the any element *e* ∈ *E* is a non-empty subset of *V*. The edges in hypergraphs are referred to as hyperedges and could connect any number of nodes. Additionally, vertices may have labels defined by a node labeling function *f*_*V*_ : *V* → Σ which is a map from the vertex set to a finite alphabet Σ. Similarly, an edge labeling function *f*_*E*_ : *E* → Ξ maps the hyperedges to a finite set Ξ of labels. A hypergraph with both vertex labels and edges labels is referred as a fully labeled hypergraph. Finally, a hypergraphlet is small (up to 4 nodes), simple, connected hypergraph. An *n*-hypergraphlet is a hypergraphlet of *n* nodes as shown in Appendix A (*n* = 1, 2, 3).

In this study, we consider a pocket as a fully labeled hypergraph where vertices are labeled by the physicochemical properties of amino acids and hyperedges (sets of amino acids) are labeled by the interaction types (e.g., hydrogen bond, spatial proximity or electrostatic). The vertex alphabet and edge alphabet are listed in Appendix B.

## 4 METHODOLOGY

### 4.1 Dataset

In this work, we focus on a protein function task: classification between enzymes and non-enzymes from protein structures. In particular, we collected a public and widely used dataset originally published by Dobson and Doig [2003]. This data set is comprised of 1178 proteins categorized as enzymes (691) and non-enzymes (487).

### 4.2 Identifying and representing protein pockets

Let *P* be a protein of interest of length *n*. We first predicted the binding pockets of *P* using *P2Rank* [23], a software tool for the prediction of ligand binding sites from protein structures. Let *P*_*C*_ denote the resulting set of predicted binding pockets of *P*. For each predicted pocket, 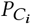, we considered five nested putative pockets 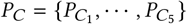such that 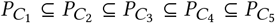. We kept the highest confidence of predicted pocket but do not label it so that the function of proteins is not leaked. This representation is unsupervised and adaptable for any downstream analysis.

Once the protein pockets *P*_*C*_ have been identified, we learned the local and global representations of these pockets. Figure 1 illustrates an example of this workflow on the bioF enzyme (8-Amino-7-oxononanoate synthase) along with corresponding protein structure (PDB ID: IDJ9) and a putative pocket (Fig. 1a). For the global representations, we start by applying discrete multiparameter persistence homology followed by multiple topological layers to produce a vectorized global representation denoted as *R*_*g*_ (Fig. 1b-c, top). In contrast, the workflow for local representations begins by creating a fully labeled hypergraph corresponding to each pocket followed by an algorithm for enumerating all *n*-hypergraphlets (1 ≤*n* ≤3). The resulting hypergraphlet-based count vectors are used as the input for a hypergraph neural network a vectorized local representation denoted as *R*_*l*_ (Fig. 1b-c, bottom). The final representation *R* is the concatenation of both local and global representations, thus, *R* = **Concat**(*R*_*l*_, *R*_*g*_).

**Figure 1:**
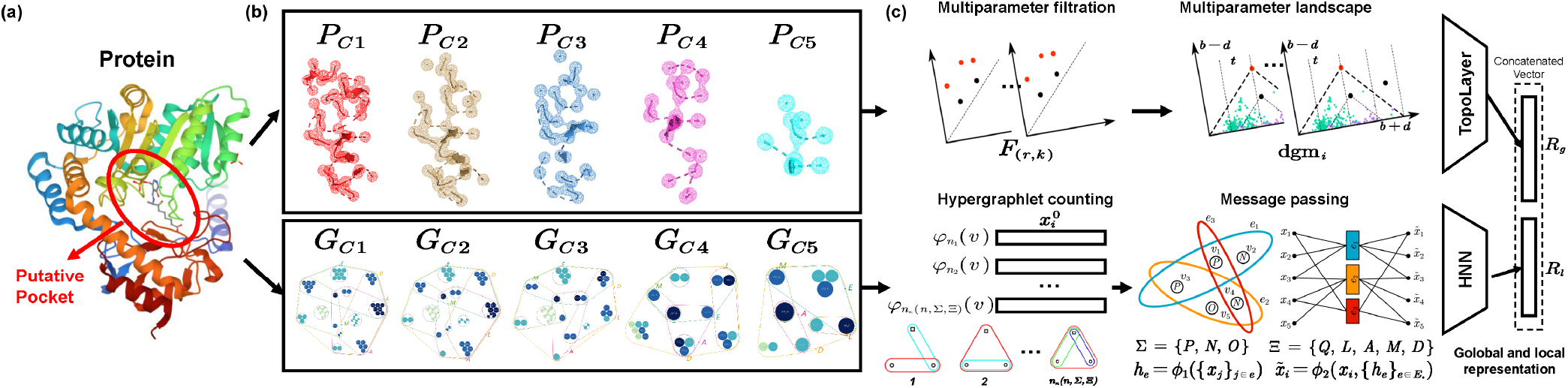
Global and local representation of a putative pocket for AONS of E.coli. (a) AONS (Gene: bioF, PDB ID: 1DJ9) is an enzyme that catalyzes decarboxylative condensation of pimeloyl-CoA and l-alanine to produce AON. (b) A *P2Rank* score filtration shows alpha carbon atoms (*C*_*α*_) around the putative pocket (top). The corresponding fully labeled hypergraphs (bottom). (c), A discrete multiparameter persistence homology and topological layers are applied to get the vectorized global representation (top). Counting of hypergraphlets (one-hot embedding is a special case) is the input of hypergraph neural network in which the local information passing is via hyperedges (bottom). The resulting concatenated vector contains both global and local information without any supervision.

### 4.3 Learning pocket geometry on a hypergraph

Typically, protein structures are modeled as protein contact vertex-labeled graphs, *G* = (*V, E*), where each amino acid residue is represented as a vertex and edges correspond to spatially close residues according to some predefined distance threshold. In this work, we enriched the graph-based representation of protein structures as follows: We first defined an edge labeling function *f*_*E*_ : *E* → Ξ such that for each edge *e* ∈ *E*, an edge label *f*_*E*_ (*e*) is introduced to represent the type of bond or interaction between the residues (e.g., hydrogen bond or disulfide bridge). The full mapping of amino acid bonds and interactions to their corresponding edge label is listed in the Appendix B. More importantly, we further generalized the representation of protein structures into fully labeled hypergraphs. For example, a proximity based hyperedge *e* is comprised of all residues within a user-defined distance threshold. This vertex- and edge-labeled hypergraph model should allow for a much better sensitivity in detecting structural templates than the previous protein structure models.

Given a fully labeled hypergraph*G* = (*V, E*) along with *f*_*V*_, *f*_*E*_, Σ, Ξ, the hypergraphlet count vector is defined as

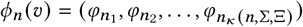

where 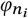 is the count of the *i*th fully labeled *n*-hypergraphlet rooted at *v* and 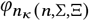 is the total number of vertex- and hyperedge-labeled *n*-hypergraphlets [26]. Given *n*, Σ and 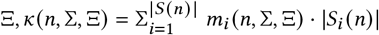 listed in Appendix B.

The count vector for each vertex is then normalized and fed into a message-passing HGNN as the initial embedding of vertices 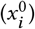. As the baseline, we also feed the HGNN with the one-hot embeddings of 20 amino acids. The message passing process in a HGNN [21] is as follows,

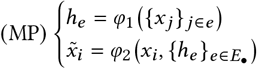

where both *φ*_1_ and *φ*_2_ are permutation-invariant functions and aggregate information from vertices and hyperedges. The exact form we used is the hypergraph equivalent to GCNII, a powerful convolutional approach with initial residual connection and identity mapping mechanisms [11].

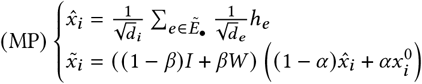

where *α* and *β* are hyperparameters, *I* is identity matrix, 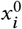 is the initial embedding of vertex 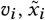 is the output embedding of vertex *v*_*i*_ after one round of message passing, *d*_*i*_ is the number of extended neighbor nodes of *v*_*i*_, *h*_*e*_ is the hyperedge embedding, *d*_*e*_ is the average degree of a hyperedge and 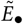 is the set of extended edges as originally described by Huang & Yang [2021].

Next, for each of the putative pockets 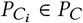, we generate a local representation 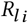. Then, the corresponding local representation for each nested putative pockets is concatenated into the final local representation for *P*_*C*_ as *R*_*l*_ = **Concat**(*R*_*l* 1_, *R*_*l* 2_, *R*_*l* 3_, *R*_*l* 4_, *R*_*l* 5_).

### 4.4 Learning pocket topology on a multiparameter perspective

The global topological representation of pockets is captured by a biparameter persistent homology, where the first filtration parameter is the distance *r* while the second filtration parameter is the *P2Rank* score *t* which measures the confident of the pocket prediction. Therefore, the sub-level set in our task is as follows

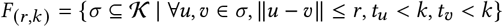

where 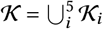 is a simplicial complex of five putative nested pockets over the metric space, (χ,*d*_χ_) of all alpha carbon (*C*_*α*_) atoms and the Euclidean distance. *u, v* represent the alpha carbon atoms with residue-wise *P2Rank* pocket score *t*_*u*_ and *t*_*v*_, respectively.

The filtration value for distance *d* ranges from 0 to the maximum diameter of all pockets by a step of 0.05 Å(1 × 10^−10^ m). The filtration value for pocket score *k* ranges from 0 to 1 but the steps are five quintiles of all residue-wise scores in the same protein. For those proteins with no or not enough putative pockets, we treat the residues between each quintile (0, 0.2, 0.4, 0.6, 0.8, 1) of *P2Rank* scores putative pockets as well. The final filtration value space is the *Cartesian product* of distance values and pocket scores. It is worth noting that we take all pockets together and use MPH to analyze the global topological information which is different to the local information capture procedure described in the previous subsection.

Next, the persistence of our biparameter filtration is computed and the persistence diagrams are obtained. The biparameter persistence diagrams are the input of a neural network with layers, Perslay, which is a unified topological layer capturing topological signatures. The unified operation towards a persistence diagram is given as

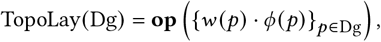

where **op**(•) is a permutation invariant operation. *w*(•) and *ϕ*(•) are the weight function and transformation function for points in persistence diagrams, respectively.

Topological signatures are automatically calculated in an topological layer with different weight and transformation functions. In this work, we focus on the topological landscape. A constant weight *w* = 1 and a *triangle point transformation ϕ*_Λ_ (Dg) = [Λ (*z*_1_), Λ (*z*_2_), …, Λ (*z*_*n*_)^⊤^. Λ (•) = max{ 0, *y* − |*z* −*x*|} is a peak function at (*x, y*). All the *z* are the regions where we want to see the landscape. The *k*th order persistent landscape could be extracted by **op** = *k*th max. Finally, for each persistence diagram Dg, we pass it to such a layer and get the global representation *R*_*g*_. Given that the second filtration value *k* is discrete, a decomposition is feasible to get the persistence diagram for sub-level sets at five pocket score intervals to map to the corresponding geometry. Such partial representation (*R*_*g* 1_, *R*_*g* 2_, *R*_*g* 3_, *R*_*g* 4_, *R*_*g* 5_) are paired with (*R*_*l* 1_, *R*_*l* 2_, *R*_*l* 3_, *R*_*l* 4_, *R*_*l* 5_) for further analysis.

### 4.5 Revealing niches by statistical analysis

The count vector representation provides very useful distribution of fully labeled hypergraphlets to build the hypergraph, showing the frequency of higher-order interactions. To statistically analyze the enrichment of these interactions, we compare our hypergraphlet-based counting with the configuration model proposed by Chodrow [2020]. Usually used as a null model, this configuration model builds random hypergraphs by holding constant node degree and edge dimension sequences but generate multiple configurations.

In this work, we extend the configuration model by adding both node labels from an alphabet Σ and hyperedge labels from an alphabet Ξ. In the null model, during sampling from the configuration, we randomly assign a node or edge label by its natural abundance. Taking a toy pocket with 3 residues and 2 interactions as an instance, as shown in Figure 2a. Consider the background abundance of node labels P, N, A are *r*_*p*_,*r*_N_,*r*_*A*_, and the background abundance of hyperedge labels D, A are 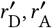, then the probability to generate such a motif is

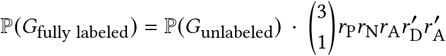

where ℙ (*G*_unlabeled_) is from the original hypergraph configuration model [12]. The equivalence classes for unlabeled hypergraphlets is shown in Appendix A.

**Figure 2:**
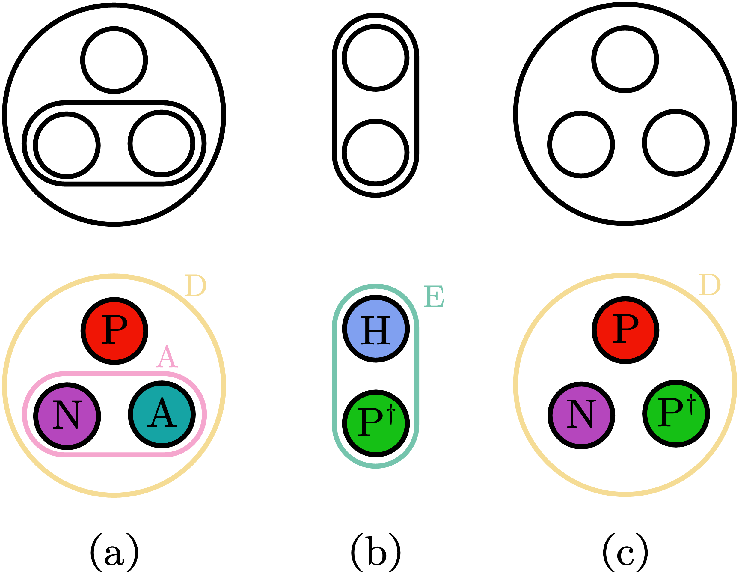
Examples of unlabeled hypergraphlets and fully labeled hypergraphlets. The string representation of each hypergraphlet is (a) *P N A*; *DA* (Type VI), (b) *H P* ^†^; *E* (Type I) and (c) *P N P* ^†^; *D* (Type II), where the top node is the root. The colors are consistent with Figure 4. (see Appendix A.1 & B.1)

We sample from our fully labeled configuration model multiple times and compute the frequency of each motif. We then acquire the over/under expression of patterns by the difference between observed count *f*_let_ and the frequency sampled from null model 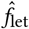. The total number of motifs are kept as the same in counting and simulation procedures. The abundance difference Δ_let_ is,

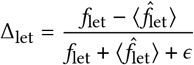

Following [27], we set the smoothing parameter ε = 4 to avoid unrealistic large values when *f*_let_ and 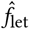 are both small.

The ensemble of over/under expression of all *n*th higher-order motifs are called hypergraph significance profile (HSP). Normalized HSP Δ_*n*_ is the fingerprint of local structure of the hypergraph [25] and has the same length as the count vector *ϕ*_*n*_ .

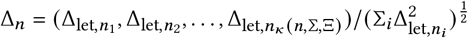

## 5 RESULTS

In this section, we evaluate both local geometric representation and global topological representation and their power for downstream analysis tasks.

For the expressiveness of our representation, we extensively evaluate how the captured geometry and topological features are aligned and consistent with experimentally verified structures in biochemistry, including mechanisms based on spectroscopic, kinetic, and crystallographic studies [35]. That is, we align the local and global information with reference pocket properties from enzymology.

### Case study of geometry

Here we give a detailed case study of the protein AONS in Figure 1. The PLP cofactor (one substrate of ANOS) is covalently bound to Lys236 while His133 and His207 are important for binding (Figure 3). Figure 4 shows that the proximity relationship is captured by a “D” hyperedge. That is, the Asp204 is hydrogen-bonded to PLP and O3 is hydrogen-bonded to His207. Such scenario is captured by another hydrogen-bond but not proximity-based labeled relationship “A” in the hypergraph. Figure 2a shows the exact hypergraphlet (*P N A*; *DA*, Type VI) representing this biochemical relationship. In the counting step, once the hypergraphlet is matched, the corresponding count will increase by one.

**Figure 3:**
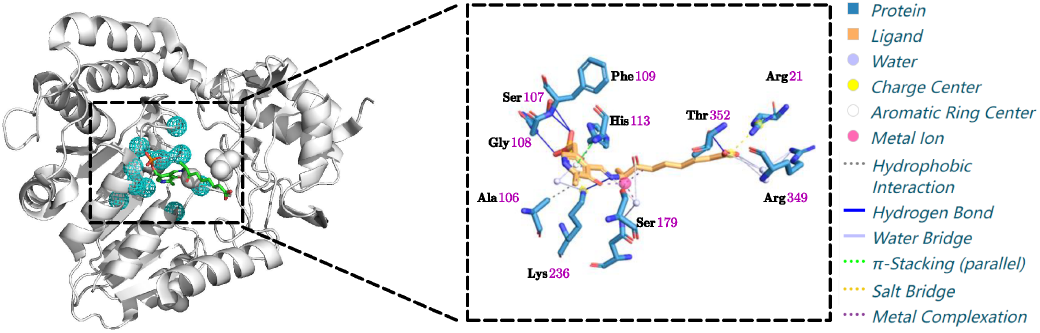
The binding pocket and interaction illustration of protein ANOS. The top 1 pocket 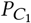 are shown as the cyan mesh. The zoom-in view of the protein, ligand and their interactions are listed at the right panel.

**Figure 4:**
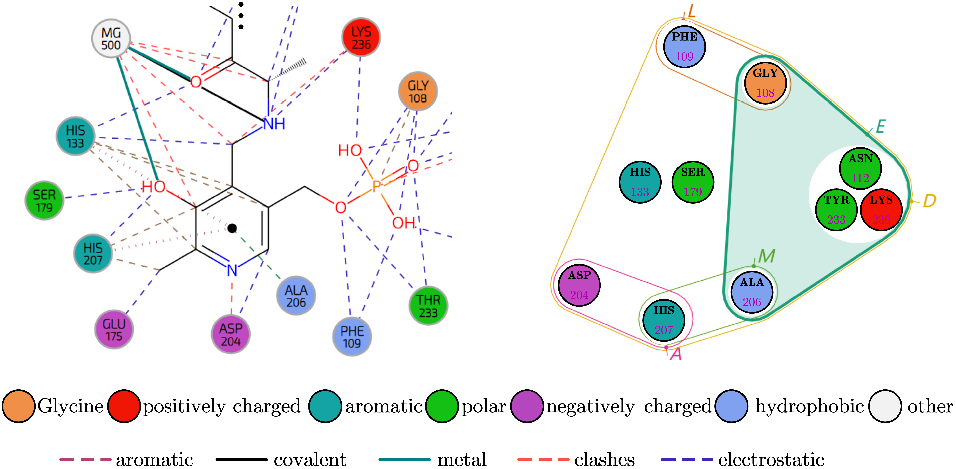
The hypergraph illustration of the top one pocket for protein ANOS. Left, part of the true 2D ligand interaction diagram. Right, the corresponding fully labeled hypergraph of the pocket.

In addition, the pyridoxal-enzyme [35] interaction can be inferred if we add the hydrophobic node label for Ala206 and Thr233 and consider the relationship “E” that captures both proximity and hydrogen bond. Figure 2b shows the corresponding hypergraphlet for this case.

However, it is not enough to just use the top one pocket. Glu175 is a negatively charged residue that can polarize the hydroxyl group of Ser179 [35]. Our top one hypergraph 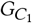 fails to capture it but Glu175 is present in 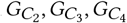 and 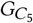. Interestingly, by considering node labels for positively (P) and negatively (N) charged residues, and polar residues (P_†_), we could identify the system of His207-Ser179-Glu175. Figure 2c shows the fully labeled 3-hypergraphlet corresponding to this configuration which is missing in 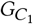 but present in 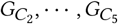 shown in Figure 4 (right). The nested pockets are necessary to dissect pockets with different sizes or pockets widely interacting with two or more functional groups.

We argue that the nested fully labeled hypergraphs are simple but enriched representations of the microenvironment within the pocket (Figure 4). Furthermore, hypergraphlets enable the incorporation of domain knowledge via the node and hyperedge labeling alphabets, thus, enabling the study of distinct complex biological interactions at the local scale.

### Case study of topology

As for the global information captured by TDA, we compared the persistent diagram for the ANOS enzyme and its five putative pockets. Figure 5 shows the Vietoris–Rips complex of ANOS (Fig. 5a) and five nested putative pockets (Fig. 5b-f) for alpha carbon atoms at maximum cutoff of *r* = 7.5Å. While most of the topological features are present, loops or voids are more prominent in these pockets.

**Figure 5:**
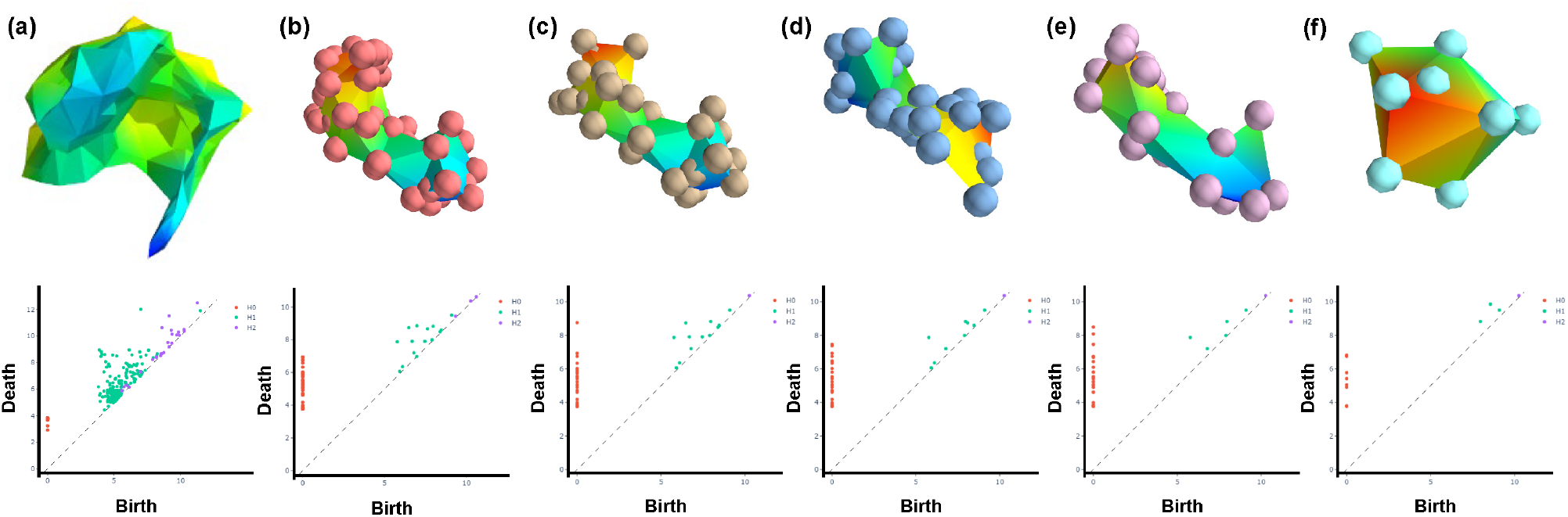
The illustration of the constructed Vietoris–Rips complex of (a) protein ANOS *P* and (b)-(f) top five putative pockets 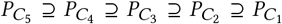. The global shape of our pockets matches the shape of our ligand (figure 3 and the top panel) quite well. MPH with nested pockets and distance (Å) shape outputs persistent diagrams (Bottom) and captures all connected components, loops, and voids with homology dimension *h* = 0, 1, 2.

The persistence diagram of ANOS’s complex is too noisy to extract pocket information. For instance, in the Vietoris–Rips complexes of the top four putative pockets, the shape is consistent to the ligand KAM, where the benzene ring and phosphonooxymethyl group are at the bottom and the keto-aminopelargonic side is at the top. In contrast, the top one pocket is smaller and only captures the rich interaction void near the benzene ring and phosphonooxymethyl group. The scattered points (*h* = 0, 1, 2) give an overview of the size and surface area of the narrow pocket. It is worth noting that the operation in the topological layer will select landscapes that correspond to the most persistent structures.

### Enzyme classification

To study the impact of learned representations, we evaluate the classification performance on the enzyme dataset [14]. Enzymes are special functional proteins speeding up the rate of a specific type of biochemical reactions. The place where the substrate binds is called the active site. Active sites are almost among the putative binding pockets [14]. Among the 1178 proteins, 691 are enzymes and 487 are non-enzymes. After removing proteins with poor structures or no pocket predictions, we keep 1139 out of 1178 proteins distributed as 666 enzymes and and 473 non-enzymes (Appendix C).

We first study the power of the sole of global or local representation. As shown in Table 1 and Figure 6, fully-labeled hypergraphlets performed better than one-hot coding of amino acids (Accuracy: 0.721 versus 0.707). More importantly, enzymes are better identified by combining local and global representations (Accuracy, Combined: 0.761; Global: 0.741; Local: 0.721), where the best performance is achieved by integrating global information to hypergraphlet-based counts. Furthermore, we study the effect of number of pockets. In Figure 6, we vary the number of pockets from top 1 to top 5 and compare the predictive accuracy. The global representation is always better than the local representation except for the top 1 pocket. However, the combined representations outperform either representation in isolation. Overall, the best accuracy for three approaches is always achieved on the top 4 or top 5 pockets.

**Table 1:** Classification accuracy for local and global representation on the enzyme dataset.

| REPRESENTATION | ACCURACY |
| --- | --- |
| GLOBAL<sup>o</sup> | 0.741±0.012 |
| LOCAL<sup>*</sup> | 0.707±0.005 |
| LOCAL<sup>*</sup>+GLOBAL<sup>o</sup> | 0.756±0.022 |
| LOCAL<sup>†</sup> | 0.721±0.021 |
| LOCAL<sup>†</sup>+GLOBAL<sup>o</sup> | **0.761±0.013** |

Mean and standard deviation of binary classification in a 10-fold cross-validation using shallow (5-layer) Neural Networks (NN).
<sup>o</sup> Topological landscape extracted by operator $k$ th max.
<sup>\*</sup> One-hot embedding as initial node features.
<sup>†</sup> Hypergraphlet counting embedding as initial node features.

**Figure 6:**
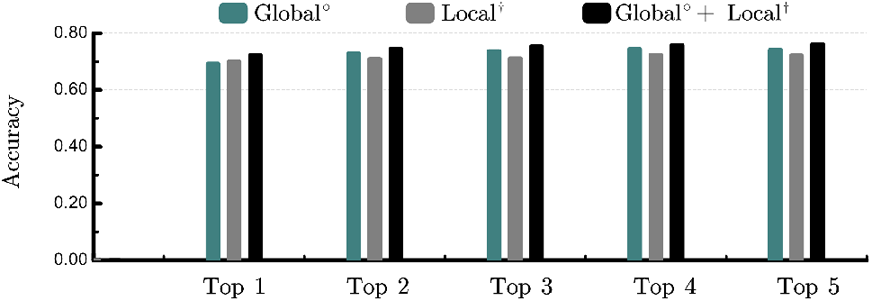
The performance comparison of different number of putative pockets. The values represent the mean accuracy based one representation Global°, Local^†^ and Local^†^ + Global°.

Then we evaluate our approaches with some start-of-the-art methods based on kernels or GNN. Since most of the work utilized one-hot encoding as node feature, we replace it with our hypergraphlet counting embedding. The addition of global features is concatenated after the final pooling or readout function. As shown in table 2, the accuracy is better with the help of global representation. The highest average accuracy 0.865 is achieved by a GNN-based approach with maximum entropy weighted independent set, SetMEWISPool [28] plus global feature. For another graph kernel-based deep learning method, DDGK[1], the best performance is achieved with one-hot embeddings instead of hypergraphlet counting. It might because the overuse of kernel tricks in both steps. We note that the global information surprisingly performs well and facilitates state-of-the-art models. These results demonstrate the advantage of our representation.

**Table 2:** Classification accuracy on multiple models.

| NODE REPRESENTATION | MODEL | w/o GLOBAL<sup>°</sup> | w/ GLOBAL<sup>°</sup> |
| --- | --- | --- | --- |
| LOCAL<sup>*</sup> | NN | 0.707±0.005 | **0.756±0.022** |
| LOCAL<sup>†</sup> | NN | 0.721±0.021 | **0.761±0.013** |
| LOCAL<sup>*</sup> | GIN(sum)[36] | 0.752±0.034 | **0.789±0.028** |
| LOCAL<sup>†</sup> | GIN(sum) | 0.773±0.021 | **0.802±0.019** |
| LOCAL<sup>*</sup> | MEWISPool[28] | 0.843±0.002 | **0.865±0.008** |
| LOCAL<sup>†</sup> | MEWISPool | 0.856±0.006 | **0.861±0.015** |
| LOCAL<sup>*</sup> | DDGK [1] | 0.831 ±0.027 | **0.853±0.019** |
| LOCAL<sup>†</sup> | DDGK | 0.827 ±0.017 | **0.848±0.012** |

Mean and standard deviation of binary classification with or without topological features. Some performance of other methods are based on
1178 proteins whereas ours is based on 1139 proteins.
<sup>°</sup> Topological landscape extracted by operator $k$ th max.
<sup>\*</sup> One-hot embedding as initial node features.
<sup>†</sup> Hypergraphlet counting embedding as initial node features.

### Statistical analysis of niches

Finally, we investigate the power of statistical analyses to associate the pocket niches with higher-order motifs [25]. We calculate the frequency and normalized HSP for fully labeled 1-, 2-, and 3-hypergraphlets in the top one pockets (Figure 7).

**Figure 7:**
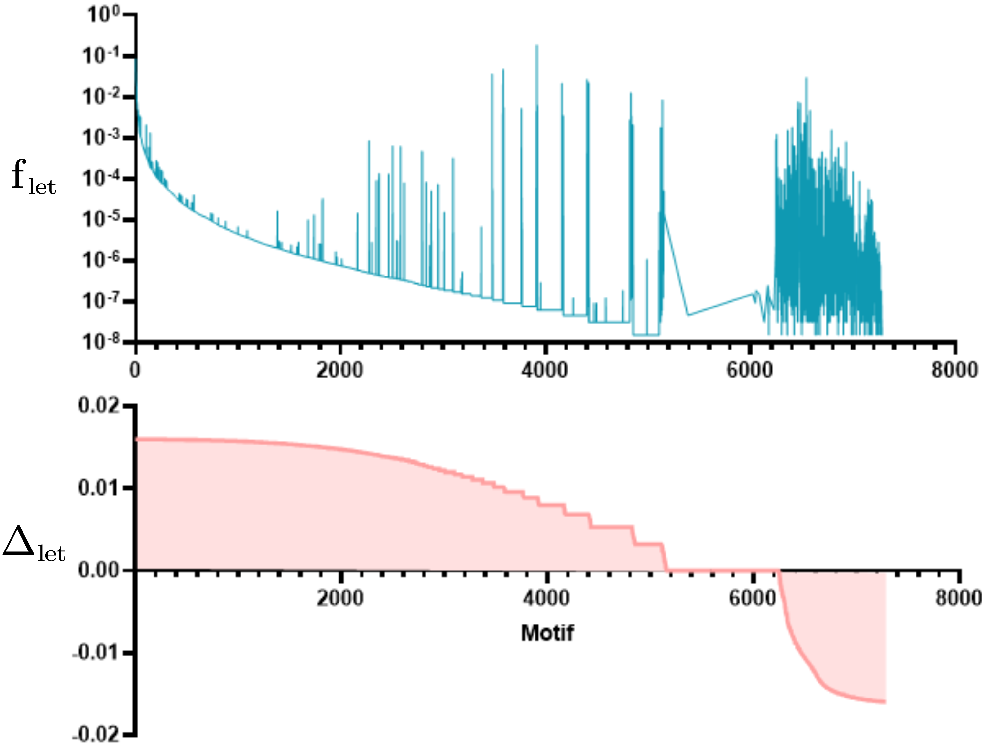
The profile of the frequency of 7, 287 fully labeled motifs including all 1-hypergraphlet, 2-hypergraphlet and 3-hypergraphlet in the scheme of positively charged (P)/negatively charged (N)/other amino acids (O). The corresponding normalized HSP Δ_*n*_ for all motifs are in descending order of Δ_let_.

Taking the charge property (Appendix B.1) as an example. We find the most abundant and significant 1-hypergraphlet, 2-hypergraphlet and 3-hypergraphlet and relate them to enzymology studies. Here we will denote a fully labeled hypergraphlet using their corresponding string representation (Appendix B.1). The most frequent and significant 1-, 2- and 3-hypergraphlets are a non-charged amino acid (“O”), two proximal non-charged amino acids (*OO*; *D*, Type I) and three proximal non-charged amino acids (*OOO*; *DD,OOO*; *DDD,OOO*; *DDDD*, Type II to X). One positively charged amino acid with two accompanying non-charged amino acids (*OOP* ; *DD*, Type IV or VI) in the pocket are immediately after in the list. Another example is a salt bridge where a glutamic acid and a lysine show an electrostatic interaction and hydrogen bond [20]. The occurrence such salt bridges could be captured by a few 3-hypergraphlets such as *P NO*; *I D* (Type III or V), *P NO*; *DI DD* (Type X) and so on. Among all 7,827 motifs identified, *P NO*; *I D* (Type III or V) is the 264th most significant hypergraphlet which is consistent with the widespread occurrence of salt bridges within proteins [20]. Unsurprisingly, our 5 physicochemical-based vertex-labeling schemes and 15 interaction types (Appendix B.2) contribute a lot for incorporating domain knowledge into downstream biological analysis tasks. The emergence of task-specific motif families could leverage the interpretability with HSP.

## 6 CONCLUSION

Locating and measuring protein pockets and cavities has been proven to be useful for biological studies. By combining TDA and GDL, we show that we can comprehensively analyze the putative protein pockets of enzymes and gain a better understanding of their niches. We present a representation learning framework for macromolecules (Figure 1), which is particularly useful when we know the structure of the underlying macromolecule. Our experimental analysis shows that learned representations encode very informative local and global feature. The extended statistical approaches for hypergraph-based motifs identified some favorable patterns in proteins structure that are consistent with enzymology. However, we note that such counting methods can become computationally expensive when we consider densely connected hypergraphs.

## A APPENDIX FIGURES

**Figure S1.**
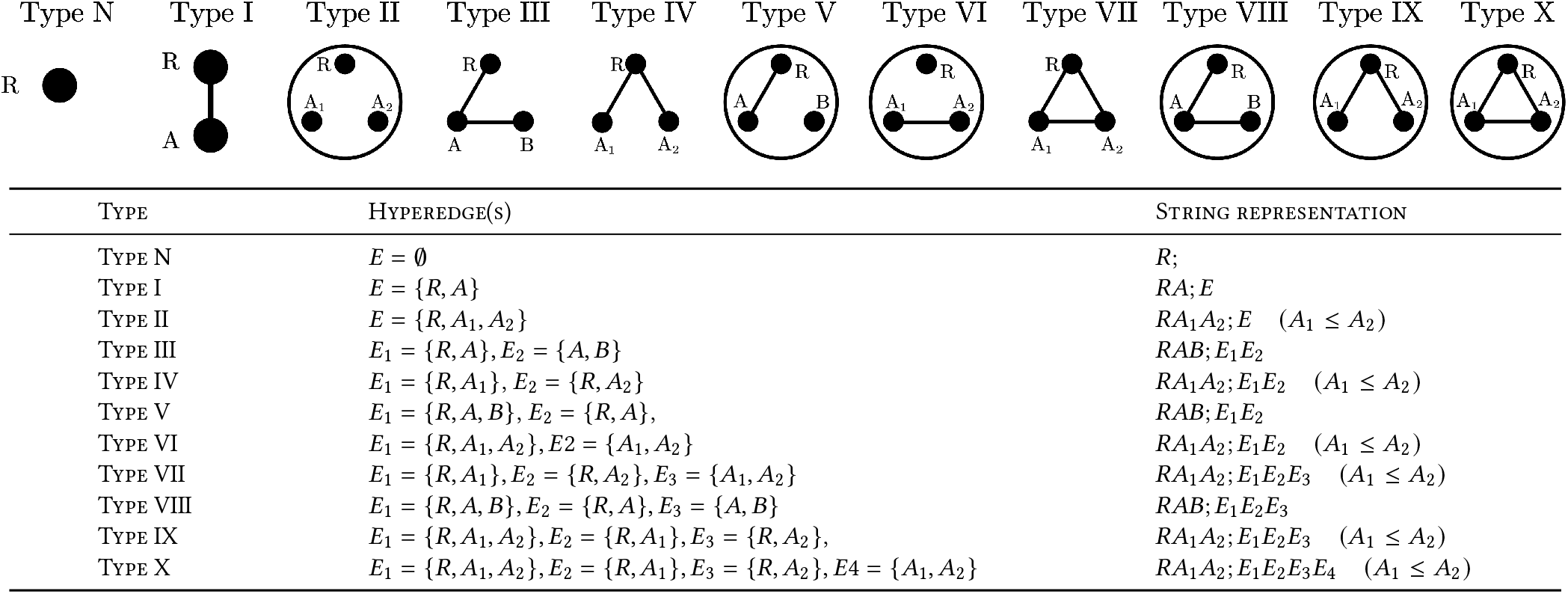
Examples of all 1-, 2- and 3-hypergraphlets without labels and their corresponding string representations.

## B APPENDIX TABLES

### B.1 Appendix tables for the vertex and hyperedge labeling alphabets

**Table S1.** The vertex labels alphabet Σ.

| NODE LABEL | AMINO ACID (SINGLE LETTER) |
| --- | --- |
| POSITIVELY CHARGED (P) | R, H, K |
| NEGATIVELY CHARGED (N) | D, E |
| OTHER (O) | A, N, C, G, Q, I, L, M, F, P, S, T, W, Y, V |
| $\vert \Sigma\vert = 3$ | |
| POLAR AND UNCHARGED SIDE CHAIN (P) | S, T, N, Q, C |
| OTHER (O) | A, R, D, G, E, H, I, L, K, M, F, P, W, Y, V |
| $\vert \Sigma\vert = 2$ | |
| SPECIAL CASE (S) | G, P |
| OTHER (O) | A, R, N, D, C, Q, E, H, I, L, K, M, F, S, T, W, Y, V |
| $\vert \Sigma\vert = 2$ | |
| HYDROPHOBIC SIDE CHAIN | A, V, I, L, M |
| OTHER | R, N, D, C, G, Q, E, H, K, F, P, S, T, W, Y |
| $\vert \Sigma\vert = 2$ | |
| AROMATIC SIDE CHAIN | F, W, Y |
| OTHER | A, R, N, D, C, G, Q, E, H, I, L, K, M, P, S, T, V |
| $\vert \Sigma\vert = 2$ | |

**Table S2.**
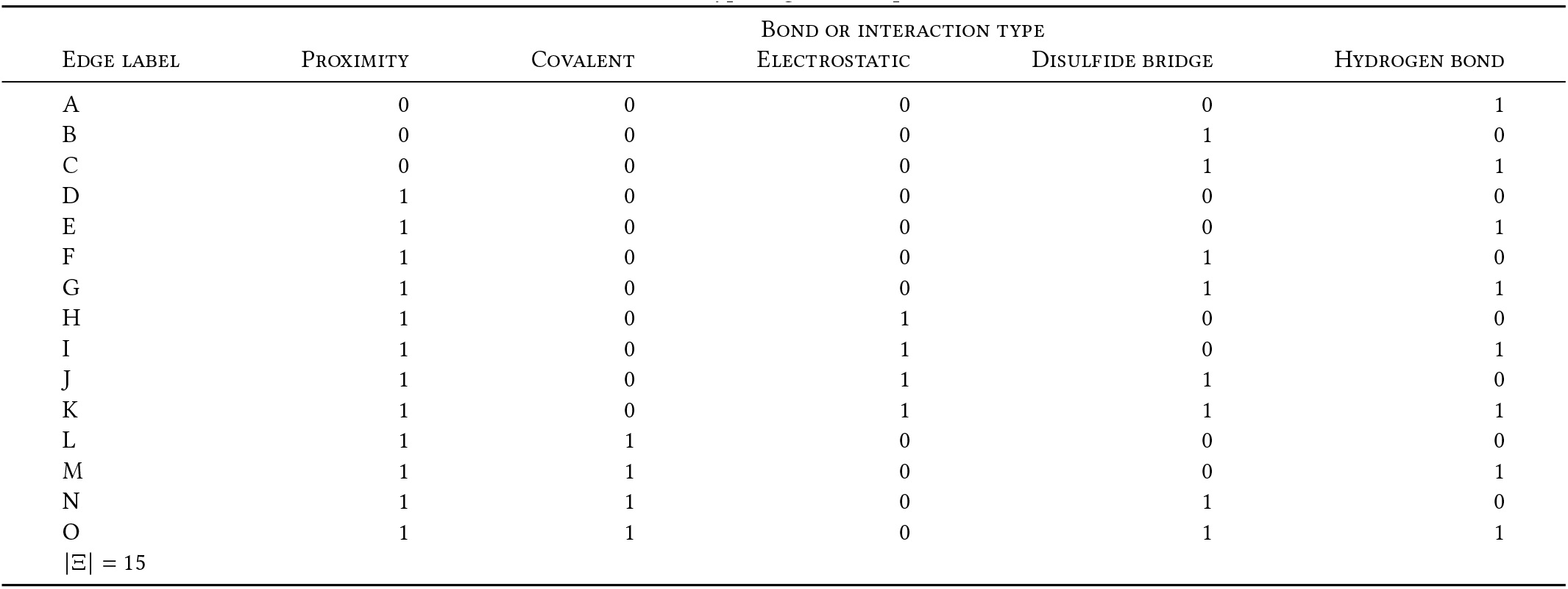
The hyperedge labels alphabet Ξ.

### B.2 Appendix table for hypergraph equivalence classes

**Table S3.**
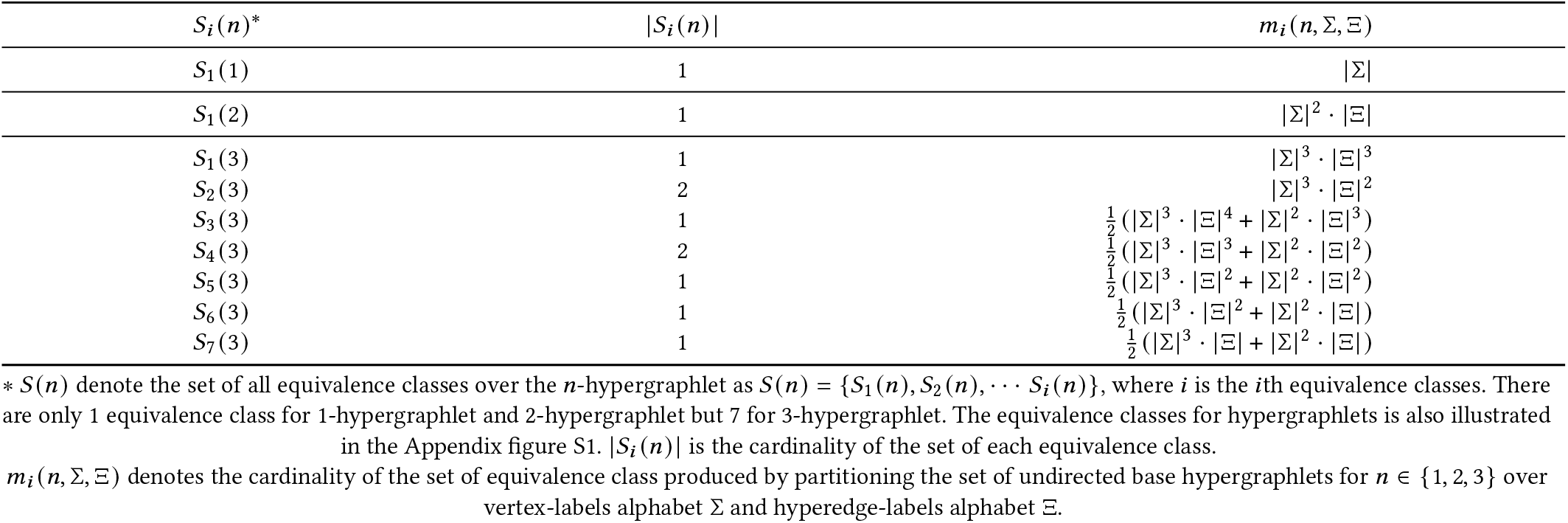
Equivalence classes over fully labeled hypergraphlets.

#### C EXPERIMENTAL SETUP

The topological representation computation is performed using rivetTDA (RIVET).

The hypergraph neural network is performed using UniGNN (UniGNN).

The topological layer is modified based on PersLay (PersLay).

The hypergraphlet counting is performed using hypergraph kernels (Hypergraphlet-kernels).

The statistical analysis is modified based on hypergraph motifs (Higher-order-motifs).

The putative pocket detection approach is *P2Rank*(P2RANK).

The enzyme dataset is originally from Dobson & Diog with 1178 proteins in which 691 are enzymes and 487 are non-enzymes (DD). We listed the Protein Data Bank (PDB) ID of 666 enzymes 473 non enzymes after filtering.

##### 666 Enzymes

11AS 1A26 1A2J 1A2P 1A33 1A49 1A4M 1A4S 1A59 1A5V 1A5Z 1A69 1A77 1A7U 1A82 1A8D 1A8P 1A8R 1A95 1A9U 1A9X 1AA6 1AA8

1ABO 1AD4 1ADE 1ADO 1AE1 1AFW 1AGJ 1AI4 1AJ5 1AJA 1AK2 1AL8 1ALN 1AQ2 1AQY 1ARC 1AUR 1AUW 1AV4 1AW9 1AY5 1AYD 1AYX

1AZ3 1AZ9 1B06 1B0E 1B0Z 1B14 1B1E 1B1Y 1B25 1B31 1B38 1B49 1B4U 1B4V 1B55 1B57 1B66 1B6A 1B6B 1B6S 1B74 1B7Y 1B80

1B92 1B9I 1B9T 1BA3 1BAI 1BC5 1BD0 1BD3 1BDO 1BGL 1BH5 1BHJ 1BIX 1BK0 1BK4 1BLI 1BMQ 1BN6 1BOX 1BS4 1BT3 1BT4 1BU6

1BUC 1BVU 1BWP 1BYS 1C02 1C1D 1C1H 1C2P 1C3F 1C3R 1C3U 1C4A 1C77 1C7I 1C7K 1C7N 1CCW 1CD5 1CEF 1CEN 1CF2 1CG6 1CIP

1CJC 1CKJ 1CL0 1CL2 1CL6 1CMV 1CNS 1CNZ 1COM 1CP2 1CPM 1CR0 1CSK 1CSM 1CVL 1CWR 1CWU 1CY0 1CYX 1D0Q 1D3H 1D4D 1D6S

1D6W 1D8I 1D8W 1DAW 1DCO 1DCU 1DD8 1DDE 1DEL 1DFA 1DHP 1DHY 1DIA 1DIH 1DIX 1DJ9 1DJN 1DJO 1DKM 1DKU 1DLJ 1DLQ 1DO8

1DQA 1DQI 1DQP 1DQX 1DS0 1DT1 1DU4 1DUC 1DUG 1DUP 1DV1 1DV7 1DVG 1DWK 1DY3 1DZT 1E0T 1E1O 1E1R 1E3D 1E3P 1E3U 1E4L

1E5L 1E5S 1E6B 1E6P 1E6U 1E7L 1E7Y 1E8Y 1E92 1EA1 1EAF 1ECJ 1EDQ 1EEX 1EG9 1EGH 1EGU 1EJ0 1EJ2 1EJB 1EKP 1EL5 1ELU

1EMV 1ENI 1EO2 1EOI 1EOM 1EOV 1EP2 1EP9 1EQC 1EQJ 1ES8 1ESD 1ESW 1EUA 1EUS 1EUU 1EVL 1EVX 1EX7 1EYB 1EYQ 1EYY 1EZ1

1EZI 1F07 1F0R 1F14 1F1J 1F28 1F2V 1F3L 1F3P 1F4L 1F52 1F5A 1F5V 1F6W 1F75 1F7T 1F83 1F8A 1F8I 1F8Y 1F9O 1F9Z 1FBT

1FC7 1FCB 1FCQ 1FG7 1FH0 1FHV 1FK8 1FO2 1FO6 1FO9 1FPX 1FR8 1FS7 1FW8 1FWK 1FWN 1FWX 1FX4 1FXJ 1FYE 1G0C 1G0H 1G0R

1G0W 1G1A 1G2O 1G3K 1G4E 1G51 1G58 1G5T 1G6L 1G6T 1G72 1G93 1G95 1G98 1G9Q 1GA4 1GA8 1GEQ 1GG1 1GGV 1GHR 1GJW 1GKX

1GMS 1GOF 1GUP 1H4V 1H6J 1H7X 1HC7 1HD7 1HDR 1HE2 1HFC 1HHS 1HI3 1HMY 1HND 1HNU 1HO4 1HP5 1HPL 1HTP 1HUC 1HW6 1HX3

1HXD 1I0S 1I12 1I1Q 1I44 1I59 1I6P 1I72 1I8A 1I9T 1I9Z 1IA1 1IAH 1IG8 1IH7 1II0 1IIP 1ILZ 1IMA 1INC 1INJ 1INO 1IO2

1IO7 1IOF 1IOW 1IRB 1ISA 1ISO 1IUS 1IVP 1J71 1J7L 1J80 1J93 1J9M 1J9Q 1JA1 1JA9 1JAE 1JAN 1JB9 1JBB 1JBP 1JBV 1JC4

1JD0 1JD3 1JDA 1JDF 1JDR 1JDX 1JEH 1JEJ 1JF9 1JFV 1JH8 1JHE 1JIN 1JK7 1JKH 1JLN 1JMF 1JN3 1JNW 1JOL 1JPR 1JQ5 1JSV

1JUK 1JWO 1JXZ 1K06 1K2O 1K89 1KAA 1KAP 1KAX 1KDN 1KEV 1KI2 1KLT 1KOQ 1KVA 1KVW 1L97 1LAM 1LCJ 1LDG 1LLO 1LOP 1LSG

1LSP 1MAC 1MAR 1MDL 1MHY 1MKA 1MLD 1MOQ 1MPP 1MUY 1NAS 1NEC 1NGS 1NHP 1NHT 1NIR 1NMT 1NNA 1NOS 1NOZ 1NSA 1OAC 1OAT

1OHJ 1OIL 1ONR 1OPR 1ORB 1OTH 1OYB 1PBK 1PGN 1PGS 1PHK 1PHM 1PHP 1PHR 1PI2 1PJC 1PLU 1PMK 1PML 1PMT 1POO 1POX 1PTR

1PUD 1PVD 1QA7 1QAE 1QAM 1QAS 1QB8 1QBB 1QBG 1QBI 1QBQ 1QCF 1QCO 1QCX 1QCZ 1QDQ 1QDR 1QF7 1QFS 1QGQ 1QH5 1QH7 1QHA

1QHH 1QHX 1QID 1QJ5 1QJC 1QJI 1QK3 1QLB 1QLT 1QM6 1QMG 1QMH 1QMV 1QNF 1QPC 1QPO 1QRE 1QRF 1QRK 1QSA 1QTR 1QVB 1R2F

1RK2 1RLR 1RLW 1RXF 1SET 1SVP 1TF4 1TKI 1TYB 1UBV 1VIF 1VNC 1VPE 1WWB 1XAA 1XAN 1XGS 1XIB 1XIK 1XKJ 1XNC 1XO1 1XPB

1XPS 1XSO 1XYS 1XZA 1YDV 1YFO 1YGE 1YGH 1YLV 1YME 1YPN 1YPP 1YTN 1ZIN 1ZRM 1ZYM 2A0B 2APS 2BSP 2CI2 2CND 2DAP 2DHN

2DIK 2DOR 2DUB 2EQL 2ER6 2ERK 2EST 2EUG 2F3G 2FGI 2FHE 2FKE 2FOK 2FUA 2FUS 2GAC 2GAR 2GD1 2GEP 2GLT 2GSQ 2HAD 2HPD

2HPR 2MAD 2MAN 2MAS 2MBR 2MIN 2MJP 2NAD 2NSY 2PF1 2PKA 2POL 2QR2 2RSL 2RUS 2SQC 2SRC 2TDT 2THF 2TLI 2TMK 2TS1 2TSC

2UBP 2UDP 2UKD 2USH 2VP3 3BIF 3BIR 3BLM 3BLS 3BTO 3CBH 3CD2 3CEV 3CGT 3CLA 3CMS 3CSC 3CSU 3CYH 3DAA 3DHE 3DMR

3ECA 3ENG 3GCB 3HAD 3PBH 3PMG 3PNP 3PRK 3PTD 3RAN 3RUB 3SIL 3SLI 3STD 3TGL 3THI 3VGC 4AIG 4DCG 4FBP 4GSA 4LZM

4MAT 4MDH 4NOS 4OTB 4PAH 4PBG 4PFK 4PGM 4TMK 5EAU 5KTQ 5YAS 6ENL 6GSS 6TAA 7AAT 7ACN 7ATJ 7CAT 7REQ 8A3H 8CHO 8LPR 9GAC

##### 473 Non-enzymes

1A04 1A0K 1A0S 1A1R 1A1X 1A21 1A2B 1A3Z 1A44 1A45 1A4X 1A62 1A64 1A7G 1A7W 1A8A 1A8O 1A92 1A99 1AAZ 1AB1 1AHQ 1AIE

1AIL 1AJJ 1ALU 1AOH 1AOX 1AQB 1AQD 1AQE 1AR0 1AS0 1AS4 1ATG 1AUE 1AUN 1AUV 1AVU 1AWP 1AXI 1AY1 1AYF 1AYI 1AYM 1B09

1B0L 1B0O 1B0Y 1B1U 1B3A 1B63 1B67 1B71 1B7D 1B7V 1B88 1B8Z 1B9N 1B9W 1BA2 1BAS 1BD8 1BFE 1BFT 1BGE 1BH2 1BHD 1BKB

1BM0 1BM3 1BMB 1BMG 1BOY 1BRX 1BTG 1BTN 1BV1 1BX8 1BXM 1BXT 1BY7 1BYF 1C1L 1C48 1C4P 1C5E 1C5K 1C6O 1C94 1CAU 1CC7

1CDH 1CDM 1CDT 1CFM 1CFW 1CI4 1CNO 1COT 1CPC 1CQ4 1CS3 1CSP 1CT5 1CXA 1CYO 1CZD 1CZQ 1D00 1D06 1D2E 1D5T 1D5W 1D7P

1DDV 1DFN 1DJ8 1DK8 1DOK 1DOT 1DQO 1DQZ 1DTJ 1DUW 1DV8 1DVN 1E00 1E0B 1E29 1E2U 1E4J 1E7C 1E7T 1E7Z 1E87 1E9L 1E9M

1EA3 1EAJ 1ED1 1EE4 1EFT 1EG4 1EGI 1EI7 1EJ4 1EKG 1EKS 1EPB 1EPU 1ERN 1ET6 1ET9 1EWF 1EYH 1EZG 1F0M 1F2L 1F2X 1F47

1F56 1F5M 1F5N 1F7C 1FAO 1FBQ 1FCY 1FD3 1FH2 1FHG 1FHW 1FL1 1FLM 1FNA 1FR9 1FSO 1FT5 1FTJ 1FVU 1FW4 1FYH 1G1C 1G33

1G3J 1G43 1G4R 1G5I 1G5Y 1G62 1G6H 1G6N 1G7C 1G7S 1G8I 1G9O 1GE8 1GPC 1GZI 1H4Y 1H75 1H8N 1H9G 1HDF 1HFA 1HG4 1HH0

1HH5 1HH8 1HJP 1HOE 1HQ3 1HQN 1HQO 1HTJ 1HUS 1HXI 1HZG 1I07 1I31 1I4U 1I6A 1I7E 1I81 1I92 1IB3 1IC0 1IC2 1IE8

1IFG 1IIT 1IKE 1IL1 1ILR 1ILS 1IN5 1IND 1INN 1INR 1IO3 1ION 1IOP 1IOZ 1IRD 1IRN 1IUZ 1IXG 1J6Z 1J73 1J7A 1J8Q

1J8S 1J8Y 1JAF 1JAH 1JB3 1JBC 1JCF 1JD1 1JDO 1JET 1JF0 1JGJ 1JI6 1JJH 1JL5 1JLJ 1JLM 1JLY 1JMW 1JOB 1JOT 1JQF

1JRR 1JVX 1JW8 1JY3 1K0K 1K33 1KDJ 1KIV 1KLO 1KMB 1KOE 1KVE 1LAF 1LE4 1LEN 1LFO 1LGH 1LIN 1LIT 1LKF 1LLA 1LOU

1LT5 1LVE 1LVK 1MDT 1MFF 1MH1 1MHO 1MJK 1MOF 1MOL 1MPC 1MRG 1MRP 1MSA 1MSC 1MUP 1MYK 1MYT 1MZM 1NAT 1NCH 1NCO

1NCX 1NDD 1NFO 1NFP 1NFT 1NG1 1NK1 1NKD 1NPS 1NSF 1NT3 1NTN 1OPC 1OR3 1ORC 1OVB 1OXY 1PCZ 1PGB 1PHN 1PLF 1PSZ

1PTX 1QAD 1QAW 1QDE 1QDN 1QDV 1QE6 1QFG 1QFV 1QG7 1QGH 1QHV 1QJ9 1QJA 1QJP 1QK8 1QKM 1QKX 1QLP 1QM7 1QME 1QOK

1QOV 1QQ1 1QQF 1QSC 1QTO 1QTP 1QVC 1R69 1RCB 1RMI 1ROP 1RPJ 1RRG 1RUX 1SCE 1SFP 1SKZ 1SWG 1UP1 1VFY 1VIN 1VPN

1WHO 1XCA 1XXA 1YFP 1YHB 1YPA 1YTT 1ZEI 1ZOP 2A2U 2ARC 2BNH 2CAV 2CBL 2ERA 2ERL 2FAL 2FB4 2FCR 2FD2 2FDN 2FGF

2FHA 2FIB 2FIT 2GAL 2GDM 2GWX 2HEX 2HIP 2HMQ 2IGD 2IMM 2IMN 2INT 2IZA 2LIG 2LIS 2LIV 2OMF 2PLH 2PSP 2TCT 2TDX

2TEP 2TGI 2TIR 2TMY 2TN4 2TNF 2TRH 2TSS 2UGI 2UTG 2VPF 2WBC 2WRP 2YGS 3CLN 3CYR 3EIP 3FIS 3GBP 3KAR 3LBD 3OVO

3POR 3PRN 3PSR 3PYP 3RHN 3SEB 3SSI 3VUB 451C 4BCL 4BJL 4ICB 4LVE 4MON 4OVO 4PAL 5PTI 7ABP 7AME 7PAZ 7PCY 7PTI 7RXN 9WGA

## D ABBREVIATION

TDA: topological data analysis
GDL: geometric deep learning
PH: persistent homology
MPH: multiparameter persistent homology
GNN: graph neural network
HGNN: hypergraphs graph neural network
PDB: protein data bank
ANOS: 8-Amino-7-oxononanoate synthase
ANO: 8-amino-7-oxononanoate
HSP: hypergraph significance profile
PLP: pyridoxal phosphate
KAM: N-[7-KETO-8-AMINOPELARGONIC ACID]-[3-HYDROXY-2-METHYL-5-PHOSPHONOOXYMETHYL-PYRIDIN-4-YL-METHANE]

## Notes

### Competing Interest Statement

The authors have declared no competing interest.

